# RNA helicase DDX5 mediates adaptive response to multi-kinase inhibitors in liver cancer

**DOI:** 10.1101/2022.03.31.486452

**Authors:** Zhili Li, Jiazeng Sun, Naimur Rahman, Bingyu Yan, Sagar Utturkar, Nadia Attalah Lanman, Majid Kazemian, Claude Caron de Fromentel, Massimo Levrero, Ourania Andrisani

## Abstract

**Objective:** To determine the role of RNA helicase DDX5 in sorafenib/multi-tyrosine kinase inhibitor (mTKI) response. Sorafenib and mTKIs downregulate DDX5 *in vitro* and preclinical hepatocellular carcinoma (HCC) models. In turn, sorafenib-mediated DDX5 downregulation activates Wnt/β-catenin and non-canonical NF-κB signaling, resulting in ferroptosis escape and mTKI resistance.

**Design and Results:** Molecular, pharmacologic and bioinformatic approaches were employed in human HCC cell lines, preclinical HCC models, and HCCs from TCGA. Earlier studies linked sorafenib effectiveness to ferroptosis. Herein we demonstrate sorafenib/mTKIs downregulate DDX5 *in vitro* and *in vivo*. To understand the effect of DDX5 downregulation, we compared TCGA-derived HCCs expressing low vs. high *DDX5* focusing on ferroptosis-related genes. *Glutathione Peroxidase 4* (*GPX4)*, a key ferroptosis regulator, was significantly overexpressed in *DDX5*^*LOW*^ HCCs. Importantly, DDX5-knockdown (DDX5^KD^) HCC cell lines lacked lipid peroxidation by GPX4 inhibition, indicating DDX5 downregulation suppresses ferroptosis. RNAseq of wild type *vs*. DDX5^KD^ cells untreated or treated with sorafenib, identified a unique set of genes repressed by DDX5 and upregulated by sorafenib. This set significantly overlaps genes from Wnt/β-catenin and non-canonical NF-κB pathways, including *NF-κB inducing Kinase* required for non-canonical NF-κB activation. Pharmacologic inhibition of these pathways in combination with sorafenib reduced DDX5^KD^ cell viability. Mechanistically, sorafenib-mediated *NIK* expression induced *NRF2* transcription, while DDX5^KD^ extended NRF2 half/life by stabilizing p62/SQSTM1, enhancing *GPX4* expression and ferroptosis escape.

**Conclusion:** Sorafenib/mTKI-mediated DDX5 downregulation results in adaptive mTKI resistance by enhancing NRF2 expression, leading to ferroptosis escape. We propose inhibition of the pathways leading to NRF2 expression will enhance the therapeutic effectiveness of sorafenib/mTKIs.

- **What is already known on this topic** – In advanced HCC, mTKIs/sorafenib, offer limited survival benefits due to resistance. Combination of VEGF (bevacizumab) and PD-L1 antibody (atezolizumab) has led to its adoption as first line treatment; however, mTKIs are still widely used in patients with contra-indications to bevacizumab; also, in atezolizumab/bevacizumab failure, mTKIs are the gold standard in second or later-line systemic therapy, emphasizing the importance of delineating the mechanism of mTKI resistance.
- **What this study adds** – We show mTKIs and sorafenib downregulate the RNA helicase DDX5. This downregulation of DDX5 by sorafenib enables activation of Wnt/β-catenin and non-canonical NF-κB pathways, leading to expression of genes that enable escape from f*erroptotic* cell death.
- **How this study might affect research, practice or policy** The Wnt/β-catenin and non-canonical NF-κB pathways activated by sorafenib/DDX5 downregulation can serve as druggable targets to enhance the anti-cancer effect of mTKIs by inducing ferroptosis, thereby opening-up new therapeutic directions against mTKI resistance.

## Introduction

Hepatocellular carcinoma (HCC) is a leading burden among all types of primary cancers with increasing incidence globally (1). Curative treatments for early stage HCC of all etiologies include surgical resection, liver transplantation and percutaneous ablation. However, high tumor recurrence rates compromise patient outcomes. In advanced HCC, multi-tyrosine kinase inhibitors (mTKIs) including systemic treatment with sorafenib (2) or lenvatinib (3), followed by regorafenib (4), cabozantinib (5), or anti-angiogenic monoclonal antibody ramucirumab (6) offer limited survival benefits due to primary or secondary resistance. Immune checkpoint inhibitors (ICIs) offer some promise (7, 8). Recently, the success of combination therapy targeting both VEGF (bevacizumab) and PD-L1 (atezolizumab) (9) has led to its adoption as a first line of treatments in many countries. However, mTKIs are still widely used in advanced HCC patients with contra- indications to ICIs and bevacizumab (10) or primary and secondary resistance associated with activated pathways for Wnt, VEGFA, and TGF-β (11, 12). Since SOR/mTKIs are still the gold standard in second or later-line systemic therapy, it is important to elucidate mechanisms of mTKI resistance, both for identifying molecular biomarkers to predict sensitivity to HCC treatment, as well as new treatment strategies to overcome resistance.

Studies by others on the mechanism of sorafenib resistance have identified crosstalk between PI3K/AKT and JAK/STAT pathways, activation of hypoxia-inducible pathways, and epithelia-mesenchymal transitions (13, 14). In addition, sorafenib treatment (15) is associated with evasion from a non-apoptotic regulated cell death called ferroptosis (16). Ferroptosis involves membrane lipid peroxidation by ferrous iron (Fe2+), under conditions of increased reactive oxygen species (ROS) (16). Importantly, CRISPR/Cas9 genome-wide screening identified Kelch-like ECH-associated protein 1 (KEAP1) as a sorafenib, lenvatinib, and regorafenib sensitivity gene in HCC (17). Upon exposure to ferroptosis-inducing compounds (e.g., erastin, sorafenib, and buthionine sulfoximine), p62/SQSTM1 inactivates KEAP1, thereby preventing degradation of the transcription factor NRF2 (15). In turn, NRF2 regulates expression of cytoprotective genes involved in glutathione production and ROS detoxification (18). However, the mechanism that regulates KEAP1 loss during sorafenib resistance has not been determined. Interestingly, earlier studies identified RNA helicase DDX5 as a key player in regulating the stability of p62/SQSTM1 (19), suggesting a role for DDX5 in sorafenib resistance.

DDX5 is a DEAD box RNA helicase (20) functioning as RNA-dependent ATPase and ATP-dependent RNA helicase *in vitro* (21). DEAD box helicases unwind RNA duplexes, displace proteins from RNA, and remodel RNA-Protein (RNP) complexes, involved in all aspects of RNA biology *in vivo* (21). DDX5 is a transcriptional regulator with critical roles in cell growth and differentiation (20), exhibiting diverse functions. In transformed hepatocytes, DDX5 regulates PRC2 function via interaction with lncRNA HOTAIR (22), STAT1 translation by resolving a G-quadruplex located at the 5’UTR of STAT1 mRNA (23), and p62/SQSTM1 degradation markedly reducing its half-life (19).

We have recently shown that liver cancer cell lines with stable DDX5 knockdown (DDX5^KD^) exhibit features of hepatic cancer stem cell (hCSC), including Wnt/β-catenin signaling activation, pluripotency gene expression, hepatosphere formation, and reduced sensitivity to sorafenib (24). Resistance to chemo/radiotherapy and molecular targeted therapies is a characteristic feature of tumors harboring CSCs (25, 26). CSCs employ various mechanisms for therapy resistance (27), including avoidance from cell death. Since DDX5 acts as a p53 co- activator (28), and sorafenib treatment results in evasion from ferroptosis (15), we investigated whether DDX5 downregulation affects sorafenib sensitivity by rescuing hepatocytes from ferroptosis.

Herein, we demonstrate that sorafenib progressively reduces expression of DDX5 in human liver cancer cell lines and in preclinical models of HCC. Comparison of the transcriptome of WT HCC cells vs. DDX5^KD^ cells as a function of sorafenib identified more than 300 common genes induced by sorafenib and repressed by DDX5. Importantly, the top-most pathways associated with those genes correspond to Wnt/beta-catenin signaling and the inflammatory, non-canonical NF-kB pathway. Wnt signaling is involved in all aspects of liver development including liver zonation, regeneration, and homeostasis (29). The non-canonical NF-κB pathway is aberrantly activated in various hematologic malignancies (30). The NF-κB-inducing kinase (NIK) encoded by the *MAP3K14* gene is a hallmark of non-canonical NF-κB activation (31), important for driving the stem cell phenotype in lung (32) and breast (33) cancers. However, the role of non-canonical NF-κB activation in HCC is unknown. Herein, we provide evidence that downregulation of DDX5 activates pathways that can serve as targets to suppress sorafenib resistance of advanced HCC.

## Materials and Methods

### Cell culture

Human HCC cell lines including wild type (WT) HepAD38^WT^ (34) and DDX5 knockdown HepAD38-DDX5^KD^ (24), Huh7, SNU387, SNU423, Hep3B, and HepaRG (35) were grown as described. Cell lines routinely tested for mycoplasma. HepAD38 cell line and its derivatives authenticated by short tandem repeat (STR) analysis performed by ATCC.

### Transfection Assays

HepAD38^WT^ (34) and DDX5 knockdown HepAD38-DDX5^KD^ cells (24) (5×10^4^ cells) transfected with TOPflash vector (100 ng) containing TCF-binding sites upstream of firefly luciferase, or NF-KB-Luciferase vector (100 ng) and co-transfected with Renilla luciferase vector (100 ng). siRNAs (50 pM) transfected using RNAiMax (Life Technologies). Luciferase activity measured 48 h after transfection using Dual Luciferase Assay system, per manufacturer’s protocol (Promega), and normalized to Renilla luciferase. Plasmids used listed in Supplementary Table S1.

### C11-BODIPY ^581/591^ assay

Cells seeded into 29 mm glass bottom dish with 14 mm micro-well #1.5 cover glass and treated with: DMSO (vehicle), sorafenib (15 μM), RSL3 (500 nM), B022 (5.0 µM), XAV939 (20 µM), ferrostatin (10 µM) or 50 pM siRNAs for 24 h, as indicated. Cells labeled with 5.0 μM C11-BODIPY ^581/591^ (Life Technologies) at 37 °C for 10 min., and visualized by fluorescence microscopy at 510 nm and 590 nm.

### Cell Viability Assays

HCC cells (1×10^4^) seeded in 96-well plates were treated with DMSO, sorafenib (7.5 μM), RSL3 (500 nM), B022 (5.0 µM), XAV939 (20 µM), ferrostatin (10 µM), or 50 pM siRNAs, as indicated, for 24 h. Growth inhibition measured at 490 nm by CellTiter 96 AQ_ueous_ One Solution Cell Proliferation assay (Promega). Viability (100 %) refers to A_490_ value of DMSO-treated cells. Background absorbance measured from wells containing media and MTS without cells.

### Huh7 xenografts

Tumor xenografts established by subcutaneous injection of 5×10^6^ Huh7 cells per NRG mouse. When tumors reached mean size of ∼180 mm^3^, mice were randomized to control and treated groups, and received vehicle (5% DMSO + 45% PEG400) or sorafenib orally at 40 mg/kg daily for first 7 days, and 80 mg/kg daily for remaining two weeks.

### HBx/c-Myc mice

Bi-transgenic HBx/c-Myc mice maintained at the Cancer Research Center of Lyon, France. Protocol authorized by the French Ministry of Education and Research (APAFIS#26268-2019121915435118 v4). Twenty week-old (4 males and 12 females) injected with Exitron nano 6000 contrast agent, and liver tumor growth monitored by micro-computerized tomography (µCT) once a week. Animals with tumor diameter of 2 mm, randomized in sorafenib-treated or vehicle group, administered sorafenib or vehicle by oral gavage five times per week. µCT monitoring continued until animal death. Liver nodules measured include those that appeared after treatment onset. Animals sacrificed at 6 weeks of treatment, or when tumor diameter was more than 12 mm (ethical euthanasia). Peri-tumor tissue and tumors were excised and frozen at - 80^0^C, or fixed in formalin.

### Immunoblotting and Immunofluorescence microscopy

methods described in detail in Supporting Information section. Antibodies employed listed in Supplementary Table S2.

### RNA preparation and qRT-PCR methods

described in detail in Supporting Information section; primer sequences listed in Supplementary Table S3, and reagents, chemical inhibitors and kits in Supplementary Table S4.

### RNA-seq analysis

HepAD38^WT^ (34) and HepAD38-DDX5^KD^ cells (24) were treated with Sorafenib (7.5 μM) for 3 days prior to harvesting. Three independent biological replicates were prepared for RNA isolation and RNA sequencing. Total RNA submitted to Novogene for quality assessment and next-generation sequencing. Paired-end 2×50 bp sequencing performed using a HiSeq2500 system (Illumina). Data quality control performed using FastQC v0.11.8. RNA expression level in each library estimated by “rsem-calculate-expression” procedure in RSEM v1.3.112 using default parameters except “--bowtie-n 1 –bowtie-m 100 –seed-length 28 --paired-end”. The bowtie index required by RSEM software generated by “rsem-prepare-reference” on all RefSeq genes, obtained from UCSC table browser on April 2017. EdgeR v3.24.013 package used to normalize gene expression and identify differentially expressed genes with following constraints: fold change > 1.5, FDR < 0.05 and TPM > 1. Gene set enrichment analysis (GSEA) performed using GSEA software (36).

### Data availability

All sequencing data are available through the NCBI Gene Expression Omnibus (GEO) database (accession number **GSE199092**).

### Statistical Analysis

Statistical analysis performed using unpaired *t* test in GraphPad Prism version 6.0 (GraphPad Software, San Diego, CA). Differences were considered significant when *p* < 0.05.

## Results

### Sorafenib downregulates DDX5 in transformed hepatocytes *in vitro* and *in vivo*

HepG2-derived HepAD38(34) cells with stable knockdown of DDX5 (DDX5^KD^) form hepatospheres when grown on low attachment plates and lack growth sensitivity to sorafenib (24). Employing HepAD38 cells as well as various other human HCC cell lines, we demonstrate that sorafenib treatment for 1-3 days progressively downregulated DDX5 (Fig. 1A-E). In addition, mTKIs regorafenib and lenvatinib also exerted a similar effect on DDX5 expression (Fig. 1F) suggesting DDX5 downregulation has a role in mTKI/sorafenib response.

**Figure 1.**
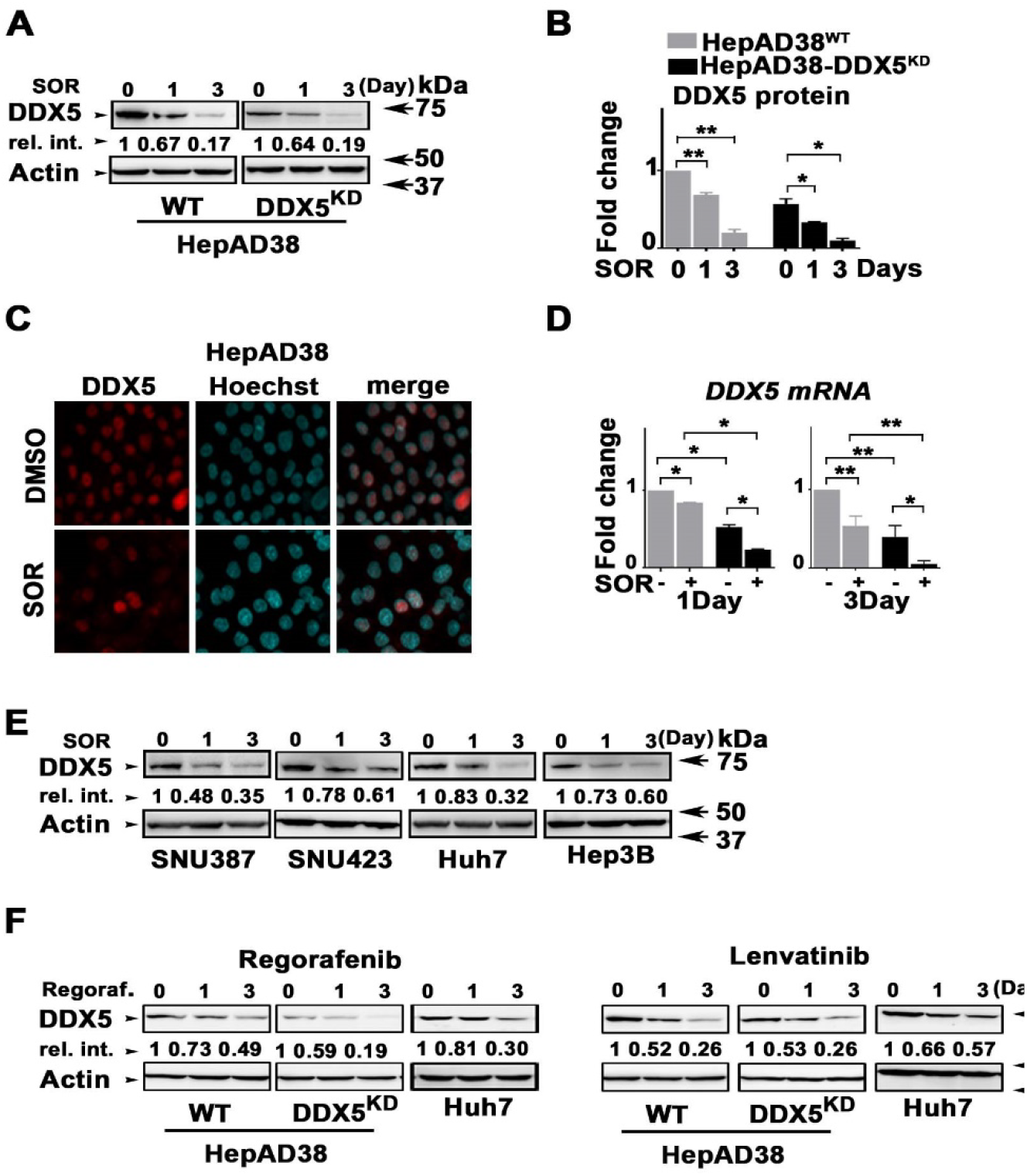
Sorafenib (SOR) downregulates DDX5. **(A)** Immunoblots of DDX5 using lysates from HepAD38^WT^ and HepAD38-DDX5^KD^ cells treated with SOR (10 µM for 1day and 7.5 µM for 3 days). **(B)** Quantification of DDX5 levels from immunoblots by ImageJ software. n=3 *p<0.05, ** p<0.01. **(C)** Immunofluorescence microscopy of DDX5 in HepAD38^WT^ cells treated with SOR (10 µM) for 1day. **(D)** RT-PCR quantification of DDX5 mRNA in indicated cell lines. n=3 *p<0.05, ** p<0.01. **(E-F)** Immunoblots of DDX5 using lysates from indicated cell lines treated with **(E)** SOR (SNU387 and SNU423: 15 µM for 1day and 10 µM for 3 days; Huh7 and Hep3B: 10 µM for 1day and 5 µM for 3 days), and **(F)** regorafenib (10 µM), lenvatinib (50 µM), as indicated. Shown, representative immunoblots from three independent experiments.

To confirm these *in vitro* observations, we used two preclinical models of HCC (Fig. 2). Huh7 xenografts in immunocompromised NRG mice and the murine HCC model of HBx/c-Myc bitransgenics (37). HBx/c-Myc bitransgenics develop liver tumors at 5-7 months without treatment with hepatocarcinogens; importantly, HBx/c-Myc HCCs (38) resemble human HCCs exhibiting a stem cell-like/progenitor phenotype (39).

**Figure 2.**
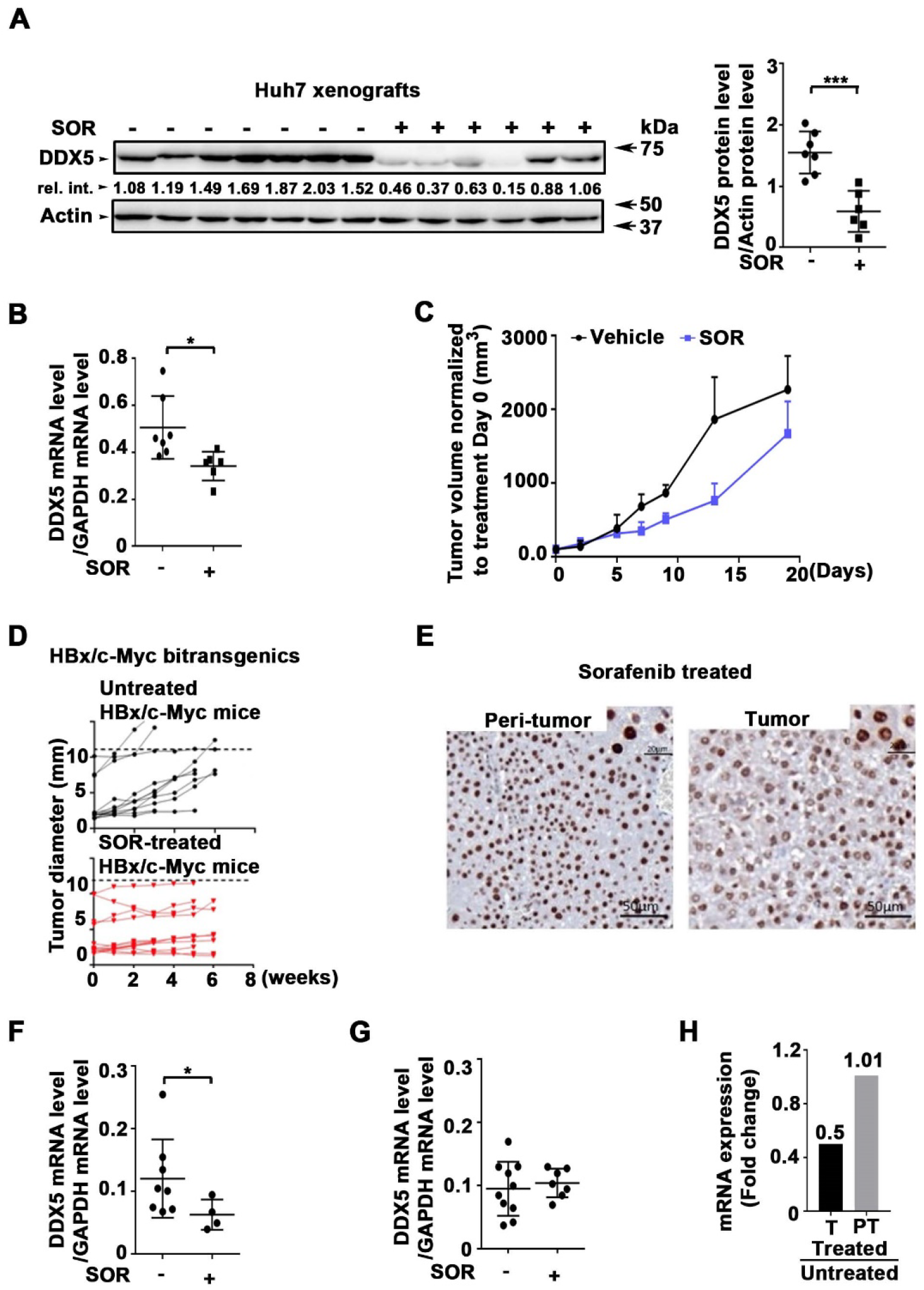
Sorafenib downregulates DDX5 in preclinical HCC models. **(A-C)** Huh7 xenografts. NRG mice bearing Huh7 tumors were treated daily with 40 mg/kg SOR for one week and 80 mg/kg SOR for two weeks (+) or DMSO (-) for 20 days. **(A)** DDX5 immunoblots from Huh7 tumors +/- SOR, as indicated. Quantification of DDX5 levels from immunoblots by ImageJ software. ***p<0.001. **(B)** Quantification of *DDX5* mRNA by qRT-PCR in tumors +/- SOR. **(C)** Tumor volume +/- SOR normalized to day-0 of treatment. **(D-H)** HBx/c-myc mice. **(D)** Tumor growth monitored by µCT scanner from each group, untreated (DMSO) and SOR-treated (60mg/kg), as indicated. **(E)** Immunohistochemistry of formalin fixed paraffin embedded (FFPE) tumor and peri- tumor stained with DDX5 antibody (Abcam ab21696, 1/1000) and counterstained with hematoxylin. **(F-H)** RT-qPCR detection of mRNA levels of *DDX5* in (**F-G**) SOR-treated *vs* untreated (DMSO) tumors and peri-tumoral tissue from HBx/c-Myc mice. *p<0.05. (**H**) *DDX5* mRNA level expressed as fold change between SOR-treated and untreated tumors and peri- tumoral tissue.

Mice bearing Huh7 tumors were treated with vehicle or sorafenib daily for 20 days; DDX5 expression in untreated vs. treated xenografts was quantified by immunoblots (Fig. 2A), and mRNA by qRT-PCR (Fig. 2B). Sorafenib significantly reduced DDX5 expression *in vivo*. Furthermore, quantification of tumor volume as a function of sorafenib suggests onset of resistance (Fig. 2C). Likewise, HBx/c-Myc mice (20 week-old) received sorafenib 5 days/week for 6 weeks. Tumor growth as a function of sorafenib (Fig. 2D) supports the HBx/c-Myc bi-transgenic mouse is a relevant model for human HCC. Sorafenib, in the HBx/c-Myc model of HCC as in Huh7 xenografts (Fig. 2C), has an effect close to that observed in HCC patients, namely, reduction of tumor growth rate, without significant tumor regression. Immunohistochemistry of DDX5 showed higher number of nuclei with “diffuse”/less intense DDX5 staining in sorafenib-treated tumors *vs*. peri-tumor (Fig. 2E), agreeing with reduced DDX5 mRNA expression in tumors (Fig. 2F, H), but not in peri-tumoral tissue (Fig. 2G, H) as a function of sorafenib.

### DDX5 downregulation is associated with ferroptosis escape

The therapeutic potential of sorafenib *in vitro* and in xenograft models of HCC is improved by pharmacologic inhibition of ferroptosis (15). Ferroptosis is a non-apoptotic, regulated cell death pathway (16). Accordingly, we examined whether downregulation of DDX5 by sorafenib has a role in ferroptosis evasion. We compared human HCCs from TCGA, those with high *DDX5* mRNA (*DDX5*^*High*^) vs. those with low (*DDX5*^*low*^) for expression of ferroptosis-relevant genes. Interestingly, *DDX5*^*low*^ HCCs revealed enhanced expression of *glutathione peroxidase 4 (GPX4)* mRNA (Fig. 3A), encoding an enzyme that suppresses lipid peroxidation, thereby rescuing cells from ferroptosis (16). Similarly, HepAD38^WT^ and HepAD38-DDX5^KD^ cells treated with sorafenib for 1-3 days, express increased *GPX4* (Fig. 3B), as well as *SLC7A11* and *NRF2*, genes involved in ferroptosis (Supplementary Fig. S1)

**Figure 3.**
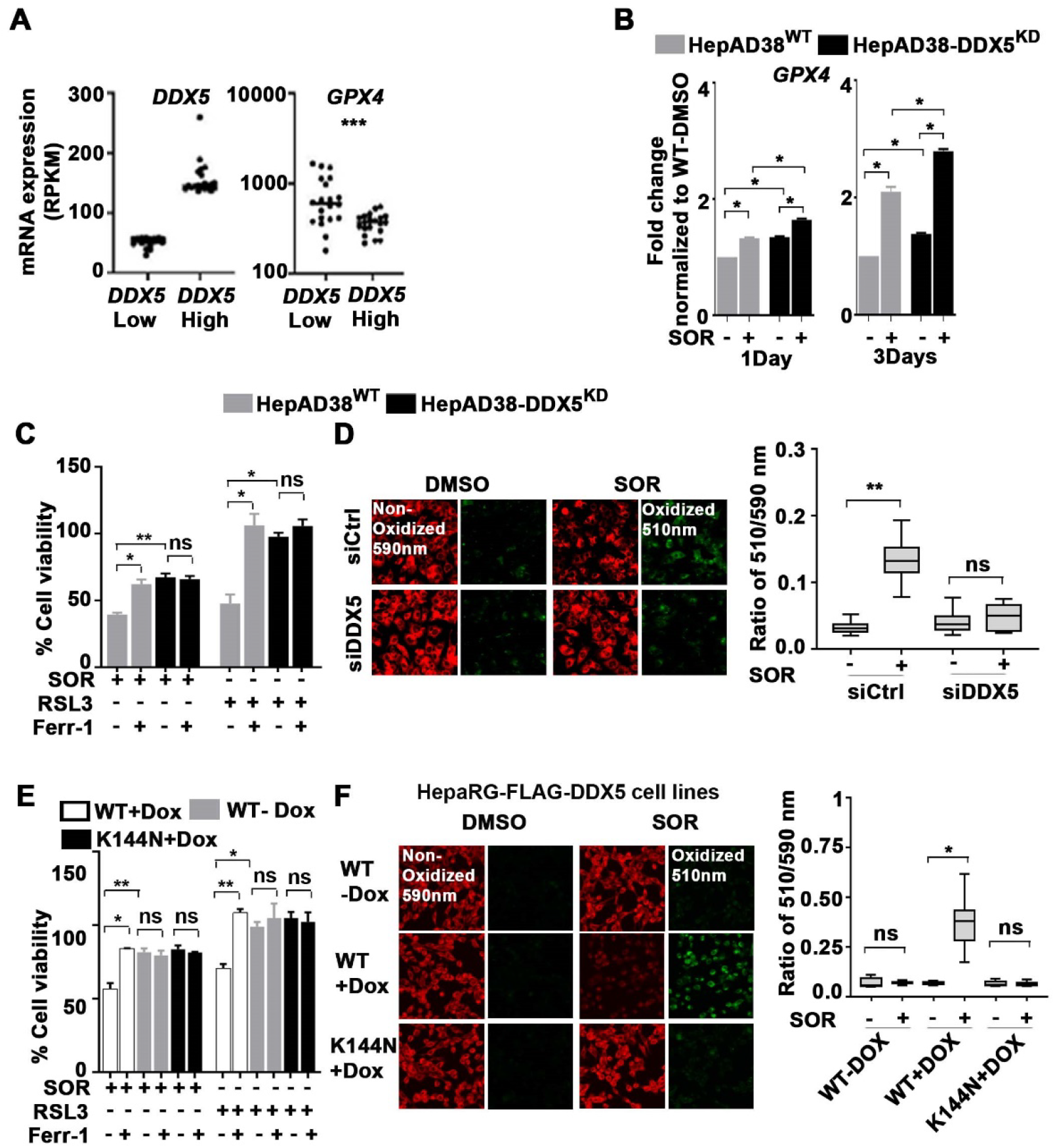
Increased expression of ferroptosis-related gene *GPX4* in HCCs with low *DDX5* expression or DDX5^KD^ cells. **(A)** HCCs from TCGA with lowest *DDX5* mRNA show increased GPX4 mRNA. Twenty HCCs analyzed per group. **(B)** qRT-PCR of *GPX4* using RNA from HepAD38^WT^ and HepAD38-DDX5^KD^ cells, treated with SOR (10 µM for 1day and 7.5 µM for 3 days, n=3). *p<0.05, ** p<0.01, ***p<0.001. **(C)** MTS viability assays of HepAD38^WT^ and HepAD38-DDX5^KD^ cells, treated with 10 µM SOR, 0.5 µM RLS3 +/- 10 µM ferrostatin-1 (Ferr- 1), as indicated, for 24 h. n=3 *p<0.05, ** p<0.01, ***p<0.00. **(D)** C11-BODIPY^581/591^ fluorescence microscopy of cells treated as indicated. (Right panel) Quantification by ImageJ software of oxidized at 510nm/non-oxidized at 590 nm. *p<0.05, ** p<0.01, ***p<0.001. **(E-F)** Doxycycline-inducible HepaRG cell lines treated with doxycycline for 24 h to express WT FLAG- DDX5 or inactive FLAG-K144N-DDX5. (**E**) MTS cell viability assays of indicated inducible HepaRG cell lines, treated with 15 µM SOR, 2.0 µM RLS3 +/- 10 µM ferrostatin-1 (Ferr-1), as indicated, for 24 h. n=3 *p<0.05, ** p<0.01, ***p<0.00. **(F)** C11-BODIPY^581/591^ fluorescence microscopy of cells treated as indicated. (Right panel) Quantification by ImageJ software. *p<0.05, ** p<0.01, ***p<0.001, (n=3).

To determine whether DDX5^KD^ cells evade ferroptosis in response to sorafenib, we performed cell viability assays as a function of treatment with ferroptosis inhibitor ferrostatin- 1(Ferr-1) or GPX4 inhibitor RSL3 (16). Sorafenib or RLS3 reduced viability of HepAD38^WT^ cells, while co-treatment with ferrostatin reversed the effect. By contrast, DDX5^KD^ cells did not exhibit any change in cell viability by RLS3 addition (Fig. 3C). Employing C11-BODIPY^581/591^, a lipid soluble fluorescent indicator of lipid oxidation (40, 41), DDX5^KD^ cells treated with sorafenib (Fig. 3D) or RSL3 (Supplementary Fig. S1C) lacked lipid peroxidation, supporting a role for DDX5 in ferroptosis. We observed the same results comparing Huh7^WT^ *vs*. Huh7-DDX5^KD^ cells (Supplementary Fig. S1D-F). The cell viability of DDX5^KD^ cells, upon treatment with sorafenib or RSL3 significantly decreased by siRNA-mediated GPX4 knockdown (Supplementary Fig. S2A). Likewise, 3-D spheroid tumor assays of HepAD38^WT^ and HepAD38-DDX5^KD^ cells, treated with sorafenib, RSL3, +/- ferrostatin exhibited the same effects (Supplementary Fig. S2B).

To determine whether the RNA helicase activity of DDX5 is required for ferroptosis, we employed doxycycline-inducible FLAG-DDX5-HepaRG cell lines (23), encoding WT-DDX5 or the inactive DDX5-K144N mutant lacking ATPase activity (42). Following doxycycline induction, overexpression of WT-DDX5 induced ferroptosis upon addition of sorafenib or RSL3, quantified by cell viability and C11-BODIPY^581/591^ assays. By contrast, the inactive DDX5-K144N mutant lacked lipid peroxidation, indicating escape from ferroptosis (Fig. 3E-F and Supplementary Fig. S2C). Taken together, these results support the enzymatic RNA helicase activity of DDX5 is required for ferroptosis, while sorafenib-mediated downregulation of DDX5 enables evasion from ferroptotic cell death.

### DDX5 regulates activation of Wnt/β-catenin and non-canonical NF-κB pathways

To determine the mechanistic significance of sorafenib-mediated downregulation of DDX5, we performed RNAseq analyses comparing the transcriptomes of HepAD38^WT^ cells and HepAD38- DDX5^KD^ cells in the presence or absence of sorafenib. Comparing DDX5^KD^ *vs*. WT cells, we identified 699 genes significantly repressed by DDX5. Using gene set enrichment analysis (GSEA), we also found that genes highly expressed in sorafenib *vs*. DMSO treated HepAD38^WT^ cells are more highly enriched in genes repressed by DDX5 (Fig. 4A). This observation suggests that a significant number of sorafenib-regulated genes are under control by DDX5. Specifically, sorafenib significantly induced 2,088 genes and DDX5 significantly repressed 699 genes, of which 313 genes are shared (Fig. 4C-D). KEGG pathway analysis of these 313 genes identified NF-κB and Wnt signaling “pathways regulating pluripotency” among the top 10 associated pathways (Fig. 4B). Regarding the “Hippo signaling pathway” (Fig. 4B), recent studies have identified the essential role of DDX5 in Hippo-mediated transcription (43). Thus, in the context of DDX5 downregulation, the Hippo pathway is transcriptionally inactive. Fig. 4D highlights select upregulated genes involved in Wnt signaling including *FZD7* and *WNT11*, while *MAP3K14, NFKB2* and *RELB* are key signaling genes of the non-canonical NF-kB pathway (31).

**Figure 4.**
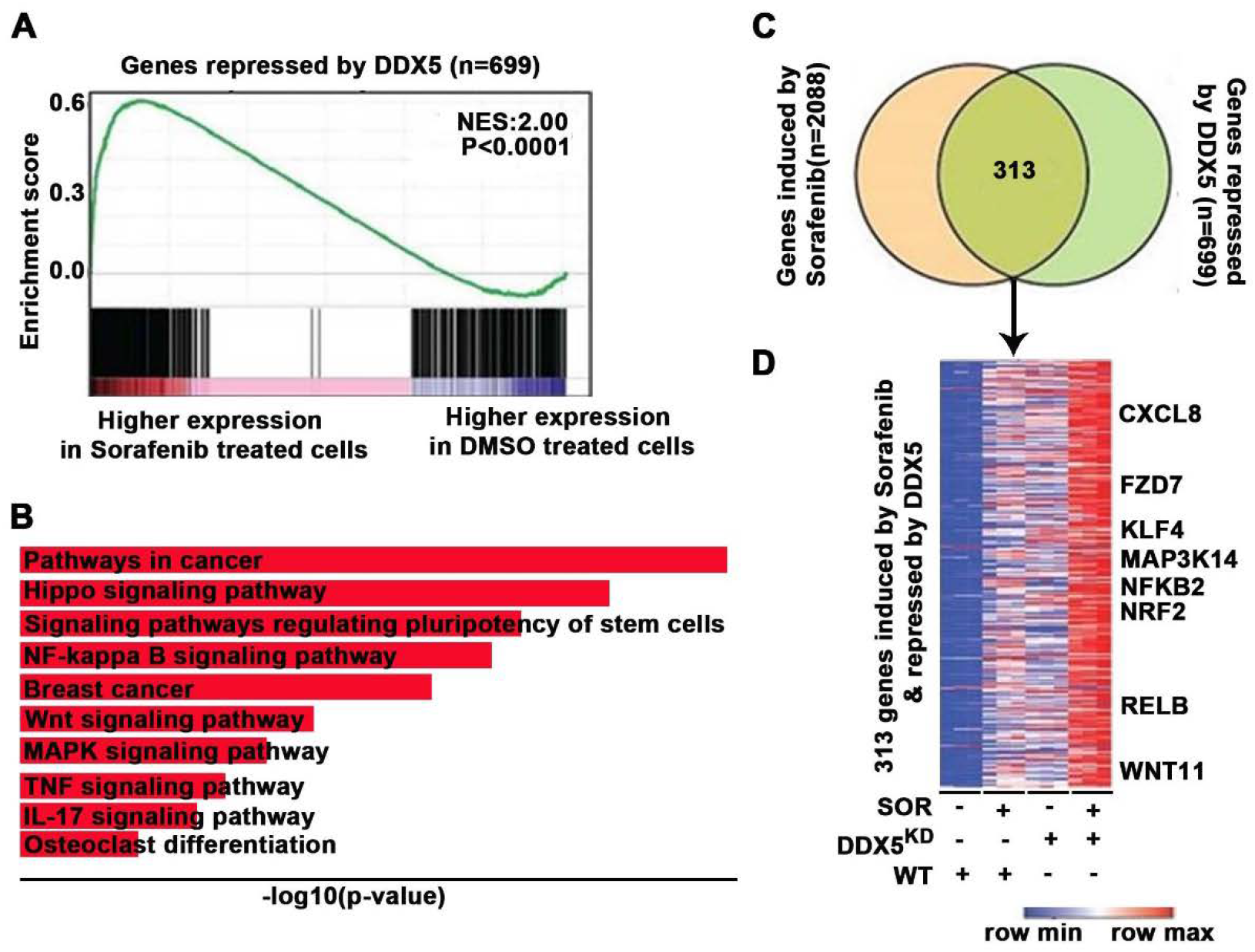
Shared sorafenib-induced and DDX5-repressed genes enriched in Wnt/β-catenin and non-canonical NF-κB signaling. **(A)** GSEA plot showing genes more highly expressed in SOR *vs*. DMSO treated HepAD38^WT^ cells, enriched in genes repressed by DDX5. **(B)** Top 10-most enriched KEGG pathways associated with genes induced by SOR and repressed by DDX5. **(C-D)** Venn diagram and heatmap of common genes between SOR-induced and DDX5-repressed genes.

### Sorafenib via DDX5 downregulation activates Wnt-β catenin and non-canonical NF-κB signaling

To determine whether sorafenib activates Wnt signaling, we performed luciferase assays using the Wnt-responsive TOPFlash reporter, containing LEF/TCF binding sites upstream of the Firefly luciferase gene. TOPFlash plasmid was co-transfected with control Renilla luciferase in various human HCC cell lines. Sorafenib increased luciferase activity, i.e., increased Wnt/β-catenin pathway activation, further enhanced by siRNA-mediated DDX5 knockdown (siDDX5), in all HCC cell lines tested (Fig. 5A). Furthermore, β-catenin immunofluorescence microscopy of HepaRG cells treated with sorafenib and siDDX5 demonstrated nuclear localization of β-catenin (Supplementary Fig. S3A). To validate these results we analyzed a commercially available PCR array comprised of more than 90 Wnt signaling genes. RNA isolated from HepAD38^WT^ cells treated with sorafenib for 3 days (Supplementary Fig. S3B) exhibited enhanced expression of Wnt signaling genes, including LRP5, DVL1, DVL3,WNT11, WNT7A, and WNT9A, confirmed by qRT-PCR (Fig. 5B).

**Figure 5.**
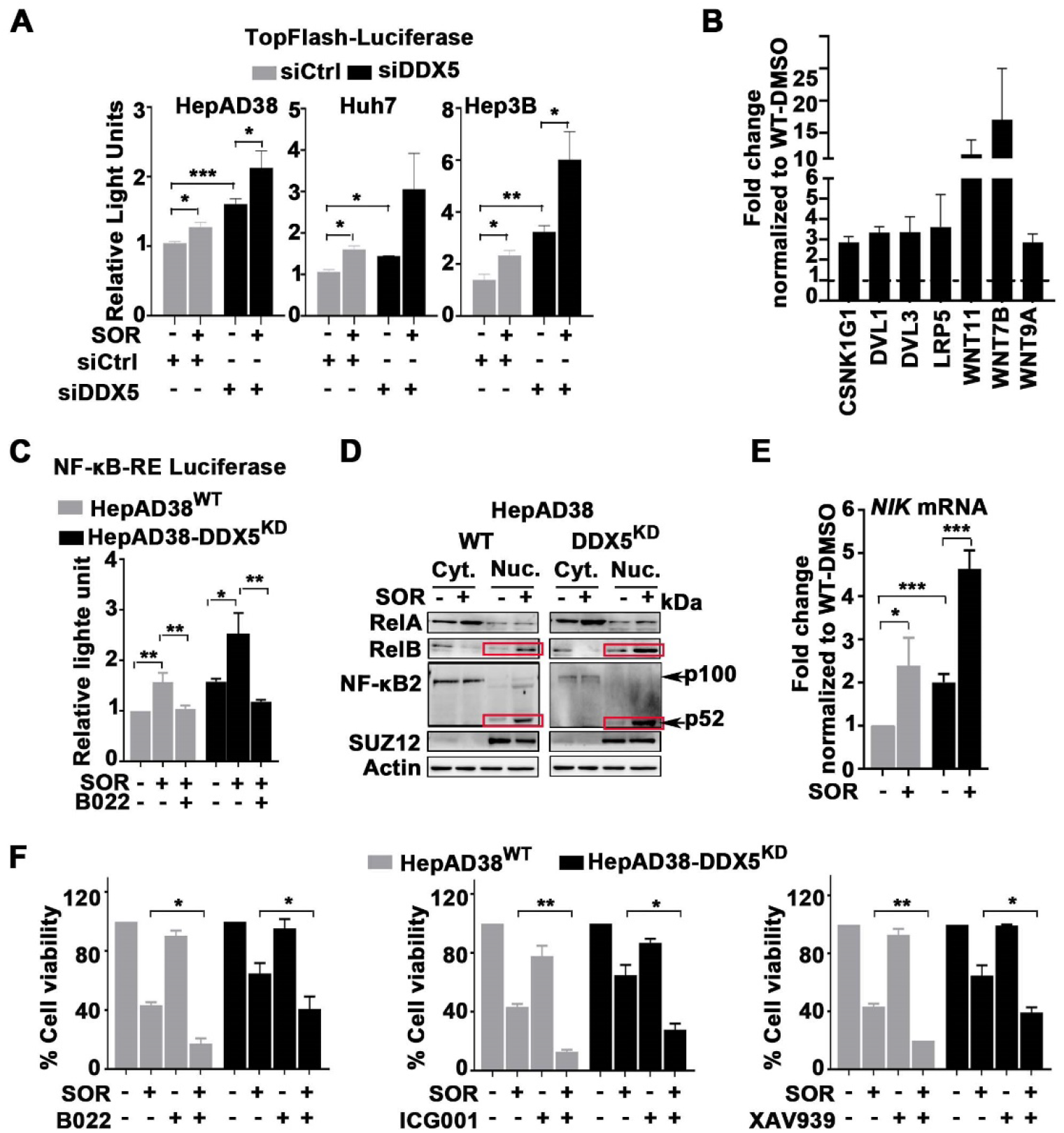
Sorafenib activates Wnt/β-catenin and non-canonical NF-κB signaling. **(A)** Wnt reporter assays using TOPFlash-Luciferase plasmid co-transfected with Renilla-Luciferase (100 ng each plasmid per 12-well plate) and siRNAs (50 pM siCtrl or siDDX5) in indicated cell lines, +/- SOR (7.5 µM). n=3 *<p.05, **p<0.01, ***p<0.001. **(B)** RT-PCR quantification of indicated Wnt regulated genes using RNA from HepAD38^WT^ cells treated with SOR (7.5 µM) for 3 days. n=3 **(C)** NF-κB reporter assays using pNL3.2.NF-κB-RE vector (Promega) +/- SOR (7.5 µM) and NF-κB-Inducing Kinase (NIK) inhibitor B022 (44) (5.0 µM), as indicated. n=3 *<p.05, **p<0.01. **(D)** Immunoblots with indicated antibodies, using cytoplasmic and nuclear lysates from HepAD38^WT^ and HepAD38-DDX5^KD^ cells +/- 7.5 µM SOR for 2 days. A representative immunoblot shown from three independent experiments. **(E)** RT-PCR quantification of *NIK* mRNA using RNA from HepAD38^WT^ and HepAD38-DDX5^KD^ cells treated with SOR (7.5 µM) for 3 days. n=3 *<p.05, ***p<0.001. **(F)** Cell viability assays of HepAD38^WT^ and HepAD38- DDX5^KD^ cells treated with SOR (10 µM), +/- B022 (5.0 µM), +/- ICG001 (15 µM) and +/- XAV939 (20 µM) for 24 h, as indicated. n=3 *<p.05, **p<0.01

To determine whether sorafenib and DDX5 downregulation activate non-canonical NF-κB signaling, we performed NF-κB-luciferase assays using HepAD38^WT^ and HepAD38-DDX5^KD^ cells as a function of treatment with NIK inhibitor B022 (44). Sorafenib increased luciferase expression in both cell lines, while B022 repressed this induction (Fig. 5C). Furthermore, sorafenib treatment of both cell lines increased levels of nuclear RelB and p52 (Fig. 5D), a hallmark of non-canonical NF-κB signaling activation (31), while it did not induce nuclear localization of RelA that mediates canonical NF-κB signaling. Sorafenib enhanced expression of NIK mRNA in both cell lines (Fig. 5E). Knockdown of NIK by siRNA transfection in HepAD38^WT^ and HepAD38- DDX5^KD^ cells inhibited sorafenib-induced processing of NFKB2/p100 to p52 (Supplementary Fig. S3C), demonstrating the requirement of NIK in non-canonical NF-κB signaling activation (31).

To determine the functional significance of these pathways in the sorafenib response, we carried out cell viability assays. Sorafenib reduced cell viability of WT and DDX5^KD^ cells by 60% and 40%, respectively. Combination treatment with sorafenib and NIK inhibitor B022, or Wnt inhibitors ICG001 (45) or XAV939 (46) further reduced cell viability in comparison to monotherapy (Fig. 5F). These results support activation of Wnt/β-catenin and non-canonical NF- κB signaling by DDX5 downregulation has a key role in sorafenib sensitivity.

### DDX5 downregulation enables transcription of *NIK* and *NRF2*

To delineate the role of DDX5 in regulating ferroptosis, we determined the effect of sorafenib on expression of *NIK*. In HepaD38^WT^ and DDX5^KD^ cells sorafenib induced *NIK* mRNA by 2-fold (Fig. 5E). Similarly, in Huh7-xenograft tumors treated with sorafenib, expression of *NIK* mRNA was elevated, as were expression of *NRF2* and *GPX4* mRNAs (Fig. 6A). In HepAD38^WT^ and HepAD38-DDX5^KD^ cells, *NRF2* and *GPX4* mRNA levels were induced by sorafenib but repressed by siRNA-mediated knockdown of *NIK* and *NFKB2*, supporting the role of non-canonical NF-κB pathway in *NRF2* transcription (Fig. 6B). In turn, knockdown of *NRF2* by siRNA transfection reduced sorafenib-mediated *GPX4* mRNA induction (Fig. 6C).

**Figure 6.**
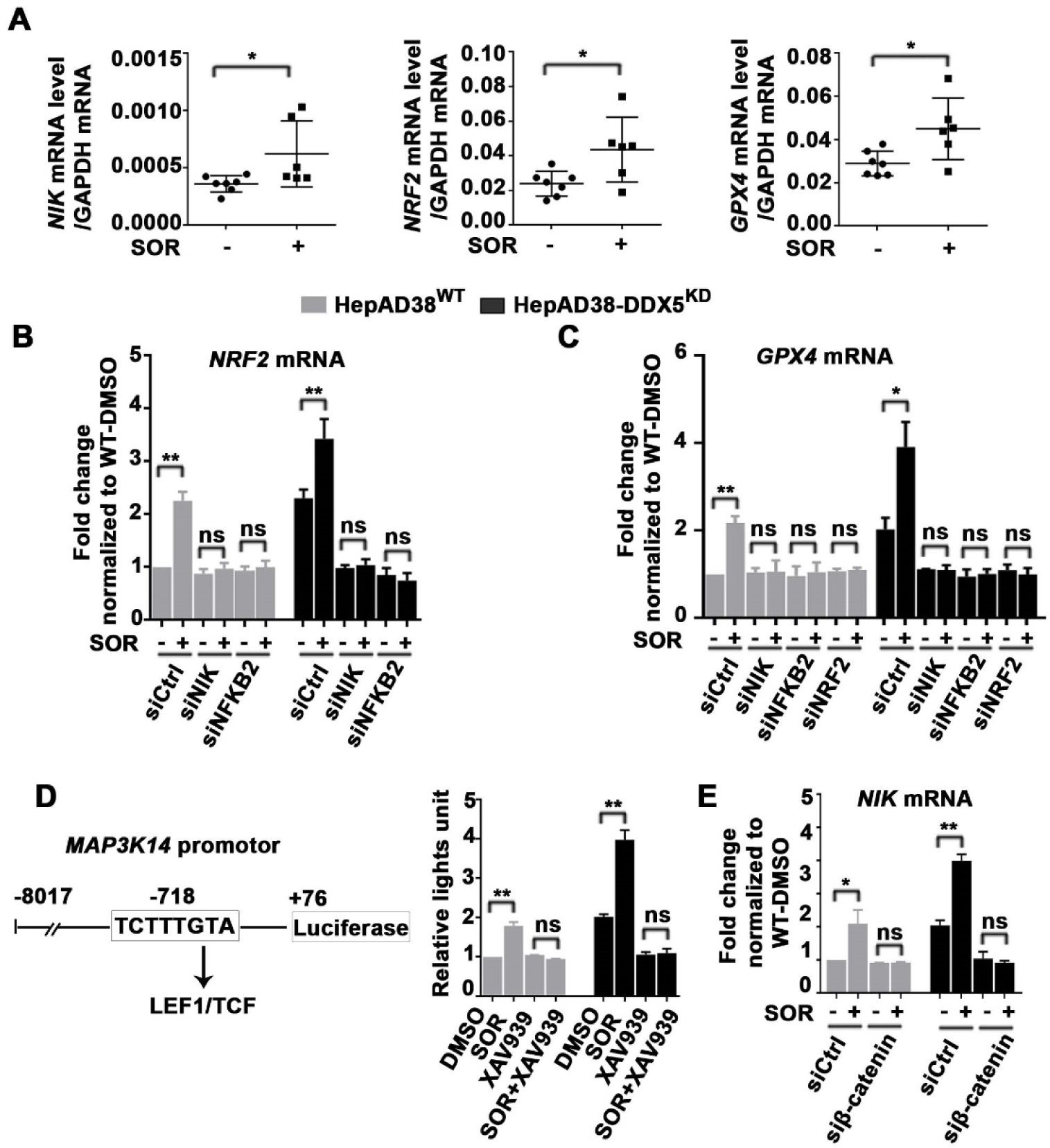
Sorafenib/DDX5^KD^ induce transcription of *NIK* and *NRF2*. **(A)** Quantification of *NIK, NRF2* and *GPX4* mRNAs by qRT-PCR in Huh7 xenograft tumors from animals treated +/- SOR for 20 days. **(B-C)** RT-PCR quantification of *NRF2* **(B)** and *GPX4* **(C)**, using RNA from HepAD38^WT^ and HepAD38-DDX5^KD^ cells treated +/- SOR (7.5 µM for 3 days) and transfected with indicated siRNAs (50 pM). n=3 *p<0.05, ** p<0.01. **(D)** *MAP3K14-*Luciferase plasmid co- transfected with Renilla-Luciferase (100 ng each plasmid per 12-well plate) in indicated cell lines, treated with +/- SOR (7.5 µM), +/- XAV939 (20 µM) or SOR+XAV939 for 48 h. n=3 *<p.05, **p<0.01, ***p<0.001. **(E)** RT-PCR quantification of *NIK* in indicated cell lines, treated +/- SOR (7.5 µM) with transfection of siRNA targeting β-catenin (50 pM) *vs*. siCtrl (50 pM).

To delineate how DDX5 regulates *NIK* transcription, we found a LEF1/TCF cis-acting element in the promoter of the *MAP3K14* gene, encoding *NIK* (Fig. 6D, left panel). Indeed, *MAP3K14-*Luciferase assays (47) displayed enhanced expression by sorafenib, inhibitable by Wnt inhibitor XAV939 (Fig. 6D, right panel). Knockdown of β-catenin by siRNA transfection inhibited *NIK* mRNA induction by sorafenib (Fig. 6E), as well as induction of *NRF2* and *GPX4* (Supplementary Fig. S4).). In summary, sorafenib via DDX5 downregulation activates Wnt signaling, which induces non-canonical NF-κB signaling and *NRF2* transcription via NIK expression.

### DDX5 regulates NRF2 protein stability

The transcription factor NRF2 coordinates expression of a vast array of cytoprotective genes under various conditions of cellular stress (18). In turn, NRF2 activity is regulated by a complex regulatory network, involving both *NRF2* transcription as well as NRF2 protein stability (18). Under normal, unstressed conditions, NRF2 half-life (t_1/2_) is approximately 20 min; disruption of the interaction between NRF2 and its potent repressor KEAP1 results in NRF2 stabilization (18). Accordingly, we determined whether sorafenib has an effect on NRF2 protein stability. In WT cells NRF2 t_1/2_ in the presence of sorafenib was longer than 45 min. Interestingly, NRF2 stability was extended in DDX5^KD^ cells, independent of sorafenib (Fig. 7A).

**Figure 7:**
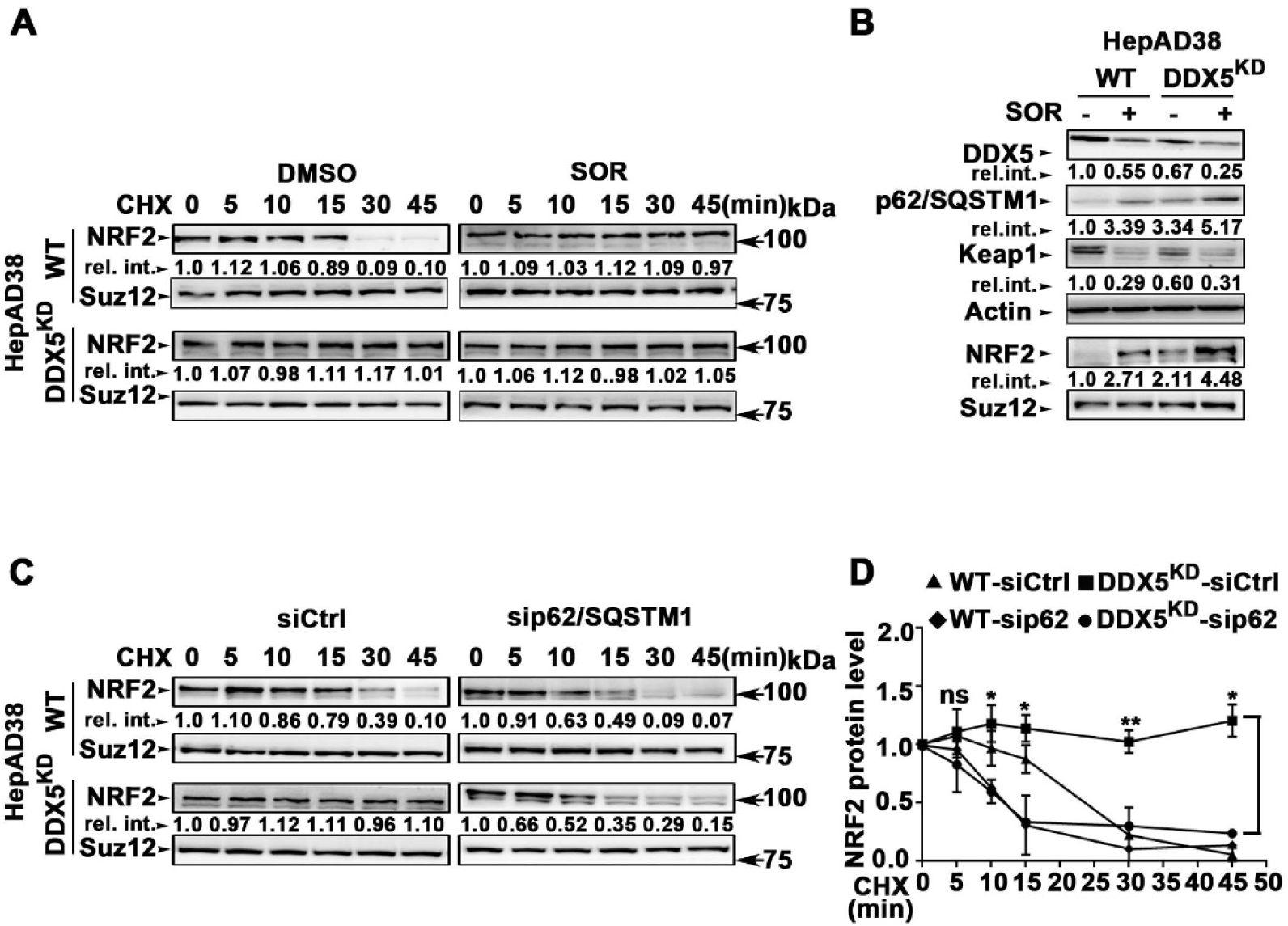
Sorafenib via DDX5 downregulation increases NRF2 half/life. **(A)** HepAD38^WT^ and HepAD38-DDX5^KD^ cells treated +/- SOR (7.5 μM for 2 days), followed by treatment with MG132 (10 μM for 4 h), extensive washing, and addition of cyclohexamide (CHX, 200 µM), as indicated. Nuclear extracts were immunoblotted with indicated antibodies. **(B)** HepAD38^WT^ and HepAD38- DDX5^KD^ cells treated with SOR (7.5 μM for 2 days). Whole cell extracts (Actin loading control) and nuclear extracts (Suz12 loading control) were immunoblotted with indicated antibodies. A representative immunoblot shown from three independent experiments. **(C)** HepAD38^WT^ and HepAD38-DDX5^KD^ cells transfected with 50 pM siRNAs (siCtrl and sip62/SQSTM1). 48 h later, cells pretreated with MG132 (10 μM for 4 h), followed by extensive washing, and addition of CHX (200 µM) for the indicated intervals. Nuclear lysates subjected to immunoblot analysis. **(D)** Quantification of half-life (t1/2) of NRF2 protein by ImageJ software of immunoblots shown in (**C**). n=3. *<p.05

A genome-wide gene editing screen identified KEAP1 as a sorafenib sensitivity gene in HCC (17). Also, earlier studies linked DDX5 to KEAP1 (19). DDX5 interacts directly with p62/SQSTM1(19), promoting p62 degradation. Thus, we reasoned, in DDX5^KD^ cells p62 is stabilized, and in turn, p62 through inactivation of KEAP1, stabilizes NRF2 (48). Indeed, immunoblots displayed increased p62 levels in DDX5^KD^ cells and upon treatment with sorafenib; conversely, protein levels of KEAP1 decreased while nuclear NRF2 protein levels increased (Fig. 7B). Next, we quantified NRF2 half-life in WT and DDX5^KD^ cells as a function of p62/SQSTM1 knockdown by siRNA and CHX addition, in a time course from 0-60 min (Fig. 7C). In DDX5^KD^ cells, NRF2 nuclear protein levels were stable longer than 45 min, similar to sorafenib treatment (Fig. 7A). By contrast, knockdown of p62/SQSTM1 in both WT and DDX5^KD^ cells reduced NRF2 t_1/2_ to approximately 10 min (Fig.7D).

In addition to this mechanism regulating NRF2 stability, lncRNA *SLC7A11-AS1* promotes gemcitabine resistance in pancreatic cancer by blocking SCF^β-TRCP^-mediated degradation of NRF2 (49). To determine if both mechanisms stabilize NRF2 in DDX5^KD^/sorafenib-treated cells, first we knocked-down by siRNAs p62 or *SLC7A11-AS1* and determined nuclear NRF2 levels, as a function of sorafenib. Knockdown of *SLC7A11-AS1* did not affect nuclear NRF2 level in the presence of sorafenib, excluding this mechanism (Supplementary Fig. S5). We conclude, DDX5 downregulation by sorafenib extends NRF2 stability through KEAP1 inactivation.

### Knockdown of NRF2 induces ferroptosis in combination with sorafenib

Inhibition of NIK by B022 in combination with sorafenib significantly reduced cell viability of both WT and DDX5^KD^ cells (Fig. 5F). To determine whether the effect of B022 on cell viability is due to ferroptosis, we assessed cell viability as a function of sorafenib in combination with NRF2 knockdown by siRNA transfection. Indeed, siNRF2 in combination with sorafenib significantly reduced cell viability of both WT and DDX5^KD^ cells, restored by ferrostatin (Fig. 8A). Employing C11-BODIPY^581/591^, we demonstrate that knockdown of DDX5 in HepAD38^WT^ suppressed lipid peroxidation in response to sorafenib; by contrast, combined knockdown of DDX5 and NRF2 by siRNA transfection induced lipid peroxidation (Fig. 8B). These results directly demonstrate the dependence on both DDX5 downregulation and NRF2 induction for ferroptosis escape. Interestingly, the absence of lipid peroxidation by DDX5 knockdown in the presence of sorafenib was abrogated by co-treatment with B022 or Wnt inhibitor XAV939, indicating both pathways have a role in ferroptosis escape (Fig. 8C). To test this hypothesis, we knocked down by siRNA transfection β-catenin or NFKB2 in combination with DDX5. Employing C11-BODIPY^581/591^, we observed enhanced lipid peroxidation in sorafenib treated cells upon combined knockdown of DDX5 and β-catenin, or DDX5 and NFKB2 (Fig. 8D). Cell viability assays performed under the same conditions, +/- ferrostatin, also support, the role of DDX5 downregulation in ferroptosis escape, via activation of non-canonical NF-κB and Wnt/β-catenin pathways (Supplementary Fig. S6A-B).

**Figure 8.**
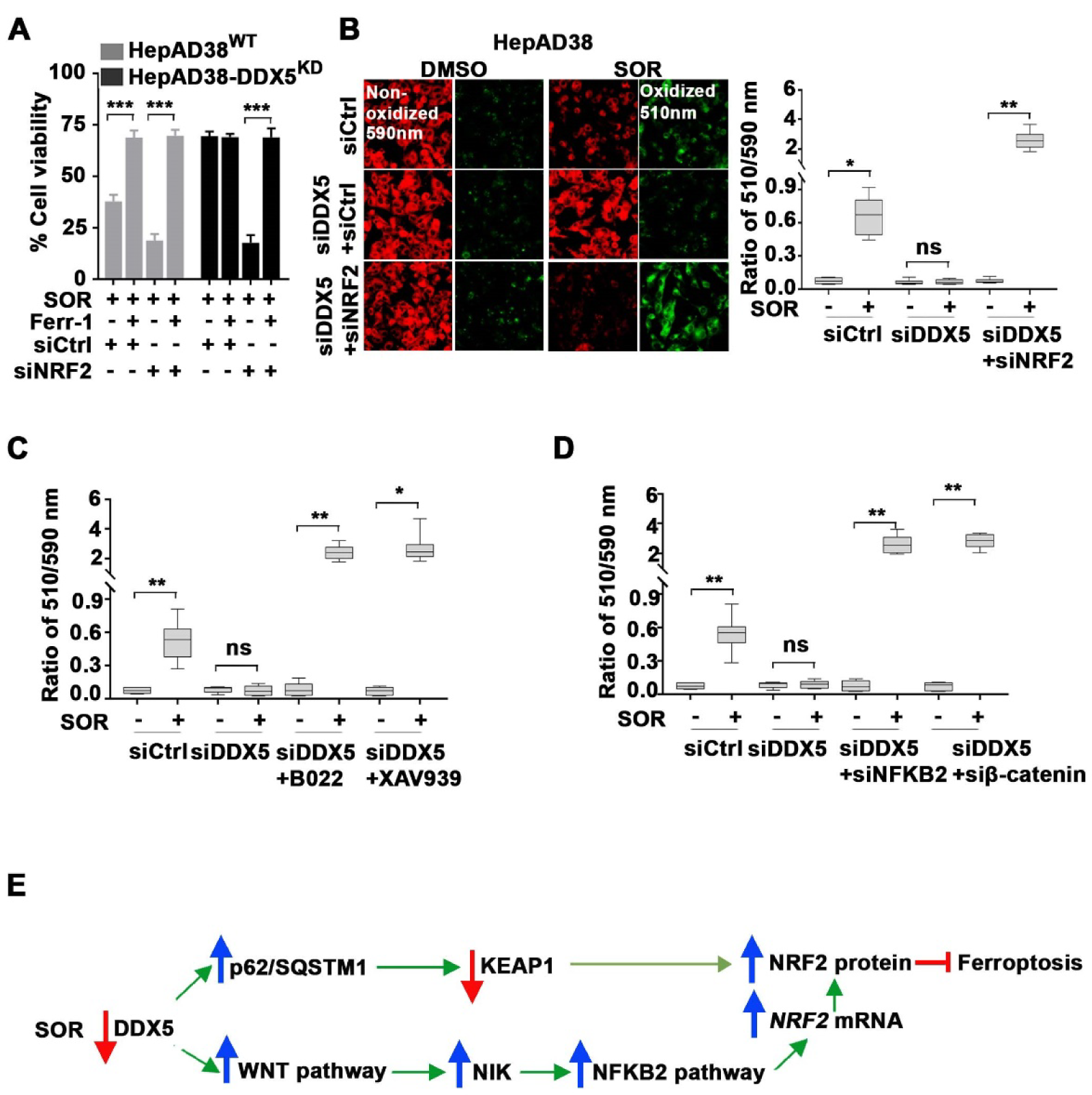
Knockdown of NRF2 induces ferroptosis in sorafenib-treated cells. **(A)** MTS cell viability assays of HepAD38^WT^ and HepAD38-DDX5^KD^ cells transfected with indicated siRNAs (50 pM siCtrl or 50 pM siNRF2) for 48 h, followed by treatment with 10 µM SOR, +/- 10 µM ferrostatin-1 (Ferr-1) for an additional 24 h. n=3 ***p<0.001. **(B)** C11-BODIPY^581/591^ fluorescence microscopy of cells treated as indicated in (A). (Right panel) Quantification by ImageJ software of 510nm/590nm ratio. n=3 *p<0.05, ** p<0.01. **(C-D)** Quantification by ImageJ of C11- BODIPY^581/591^ assays (510nm/590nm ratio) of HepAD38^WT^ cells treated +/- SOR (15 µM for 24 h) and transfected **(C)** with siCtrl (50 pM) or siDDX5 (50 pM) in combination with B022 (5.0 µM) or XAV939 (20 µM) or (**D**) with 50 pM each of siCtrl or siDDX5 in combination with siNFKB2 or siβ-catenin. n=3 *p<0.05, ** p<0.01. (**E**) Diagram of the mechanism by which sorafenib-mediated downregulation of DDX5 results in ferroptosis evasion.

## Discussion

In this study, we provide evidence supporting the role of the RNA helicase DDX5 in orchestrating mTKI/sorafenib resistance. We show that sorafenib and mTKIs downregulate expression of DDX5 in various human HCC cell lines and preclinical models of HCC (Fig. 1 and 2). Since DDX5^KD^ cell lines exhibit hepatosphere formation, reduced sensitivity to sorafenib, and increased expression of pluripotency genes, all characteristic of CSCs (24), we reasoned downregulation of DDX5 by sorafenib is a likely contributing factor to sorafenib resistance. Herein, we explored this mechanism.

### DDX5 downregulation allows ferroptosis escape

Earlier studies demonstrated an association between sorafenib sensitivity *in vitro* and *in vivo*, and pharmacologic inhibition of ferroptosis (15). Since sorafenib downregulates DDX5, using human HCCs from TCGA, we found that *DDX5*^*low*^ HCCs express statistically significant increased mRNA level for *GPX4*, encoding an enzyme that protects cells from ferroptosis (Fig. 3A). Similarly, enhanced expression of *GPX4* occurs in sorafenib treated WT cells and DDX5^KD^ cells (Fig. 3B). Cell viability assays in monolayer (Fig. 3C) or 3-D spheroid cultures (Supplementary Fig. S2B) demonstrated that GPX4 inhibition by RSL3 increased ferroptotic cell death only in WT cells, reversed by ferrostatin. Likewise, lipid peroxidation, quantified by C11-BODIPY^581/591^ as a function of sorafenib, increased in WT but not DDX5^KD^ cells (Fig. 3D). Significantly, the RNA helicase activity of DDX5 is required for ferroptosis induction (Fig. 3E and F), suggesting that functionally active DDX5 represses transcription of genes mediating rescue from ferroptosis.

### Sorafenib/DDX5 downregulation activates Wnt/β-catenin and non-canonical NF-κB signaling

Employing RNAseq analyses of WT and DDX5^KD^ cells treated with sorafenib, we identified common genes induced by sorafenib and repressed by DDX5 (Fig. 4). These include genes associated with Wnt/β-catenin and non-canonical NF-κB pathways. We employed various Wnt- and NF-κB-responsive assays and demonstrate that sorafenib/DDX5^KD^ activate both pathways (Fig. 5A-E). Importantly, combined treatment of sorafenib and NIK inhibitor B022 (44) or Wnt inhibitors, ICG001 (45) or XAV939 (46), significantly reduced cell viability of both WT and DDX5^KD^ cells (Fig. 5F). We conclude, these pathways (Wnt/β-catenin and non-canonical NF- κB) activated by DDX5 downregulation play an important role in sorafenib sensitivity.

In HCC, dysregulated activation of Wnt signaling is mediated by activating mutations of *CTNNB1* (β-catenin) and/or deregulated expression of Wnt signaling components (50, 51). Interestingly, TCGA-derived HCCs linked *DDX5*^*low*^ tumors to increased expression of DVL1 (24), an indispensable, intracellular regulator of Wnt activation (52, 53). However, further studies are needed to understand how DDX5 represses Wnt-regulated gene expression (54). Active Wnt/β- catenin signaling is associated with hCSCs (55, 56), contributing to poor prognosis and immunosuppression (11, 12). Regarding activation of non-canonical NF-κB pathway, it involves nuclear translocation of RelB and NFKB2/p100 processed to p52, which requires de novo synthesis and accumulation of the NF-κB-inducing kinase (NIK), encoded by the *MAP3K14* gene (31, 57). NIK is important for development of the CSC phenotype in various cancers (32, 33), although its role in HCC is unknown. We show that expression of *NIK* increased by sorafenib/DDX5^KD^ *in vitro* (Fig. 5E), as well as in Huh7 xenograft tumors *in vivo* (Fig. 6A). NIK, via NF-κB activation, is required for increased expression of *NRF2* and *GPX4* mRNAs in WT and DDX5^KD^ cells treated with sorafenib (Fig. 6B-C). Furthermore, *NRF2* and *GPX4* mRNA levels were also elevated in sorafenib-treated Huh7 xenografts (Fig. 6A).

### Sorafenib/DDX5 downregulation via NRF2 mediate ferroptosis escape

Various studies have identified a mechanistic connection between Wnt signaling and canonical NF-κB activation (58). However, it has been unknown whether Wnt signaling also activates non-canonical NF-κB. Herein we provide evidence for a direct relationship between Wnt/β-catenin signaling activation by sorafenib/DDX5^KD^ and transcription of the *MAP3K14* gene, a prerequisite for non-canonical NF- κB activation (Fig. 6D). In turn, active NF-κB mediates *NRF2* transcription (Fig. 6B), as also shown by others (59). Importantly, DDX5 knockdown, in addition to the transcriptional induction of *NRF2*, also regulates NRF2 half-life. In normal growth conditions, NRF2 t1/2 is approximately 15 min; sorafenib extended NRF2 half-life to longer than 45 min (Fig. 7A) via disruption of its interaction with KEAP1 (Fig. 7B). Earlier studies linked loss of DDX5 to p62/SQSTM stability, resulting in reduced KEAP1 protein levels (19). Indeed, in DDX5^KD^ cells KEAP1 levels decreased, while p62 levels increased (Fig. 7B). Knockdown of p62/SQSTM reduced NRF2 half-life to 10 min. in both WT and DDX5^KD^ cell lines (Fig. 7C-D). Thus, DDX5 downregulation has a dual role in sorafenib sensitivity, by regulating transcription and protein stability of NRF2.

Cell viability and C11-BODIPY^581/591^ assays, in the context of sorafenib/DDX5 knockdown, also demonstrate the important role of NRF2 in cell survival and ferroptosis escape, respectively (Fig. 8A-B). Firstly, both Wnt and non-canonical NF-κB signaling regulate *NRF2* transcription (Fig. 6B and Supplementary Fig. S4A). Secondly, lipid peroxidation/ferroptosis induction occurs by pharmacologic inhibition of NIK or Wnt/β-catenin signaling (Fig. 8C), or by knockdown of β-catenin or NFKB2 (Fig. 8D). These results demonstrate a mechanistic connection between Wnt/β-catenin and non-canonical NF-κB signaling in ferroptosis escape, as shown in the diagram of Fig. 8E, and identify both pathways as druggable targets for enhancing the anti-cancer effectiveness of mTKIs via induction of ferroptosis.

## Acknowledgements

The authors thank P-PAC animal facilities and imaging platform at Cancer Research Center of Lyon, in particular Thomas Barré and Emile Servoz for mouse treatment and follow-up.

