## Supplementary information for "RNA helicase DDX5 mediates adaptive response to multi-kinase inhibitors in liver cancer"

[1]Department of Basic Medical Sciences, [2]Purdue Center for Cancer Research, [3]Department of Biochemistry, [4]Department of Comparative Pathobiology, [5]Department of Computer Science, Purdue University, West Lafayette, IN 47907, USA, [6] Cancer Research Center of Lyon (CRCL) - INSERM U1052, CNRS5286, Univ Lyon, Université Claude Bernard Lyon 1, F69000 Lyon, France, [7]Hospices Civils de Lyon, Service d'Hépatologie et Gastroentérologie, Groupement Hospitalier Lyon Nord, France.

Department of Basic Medical Sciences,
Purdue University
201 S. University Street
West Lafayette, IN 47907-2064

**Supplementary information:**

**Figures**

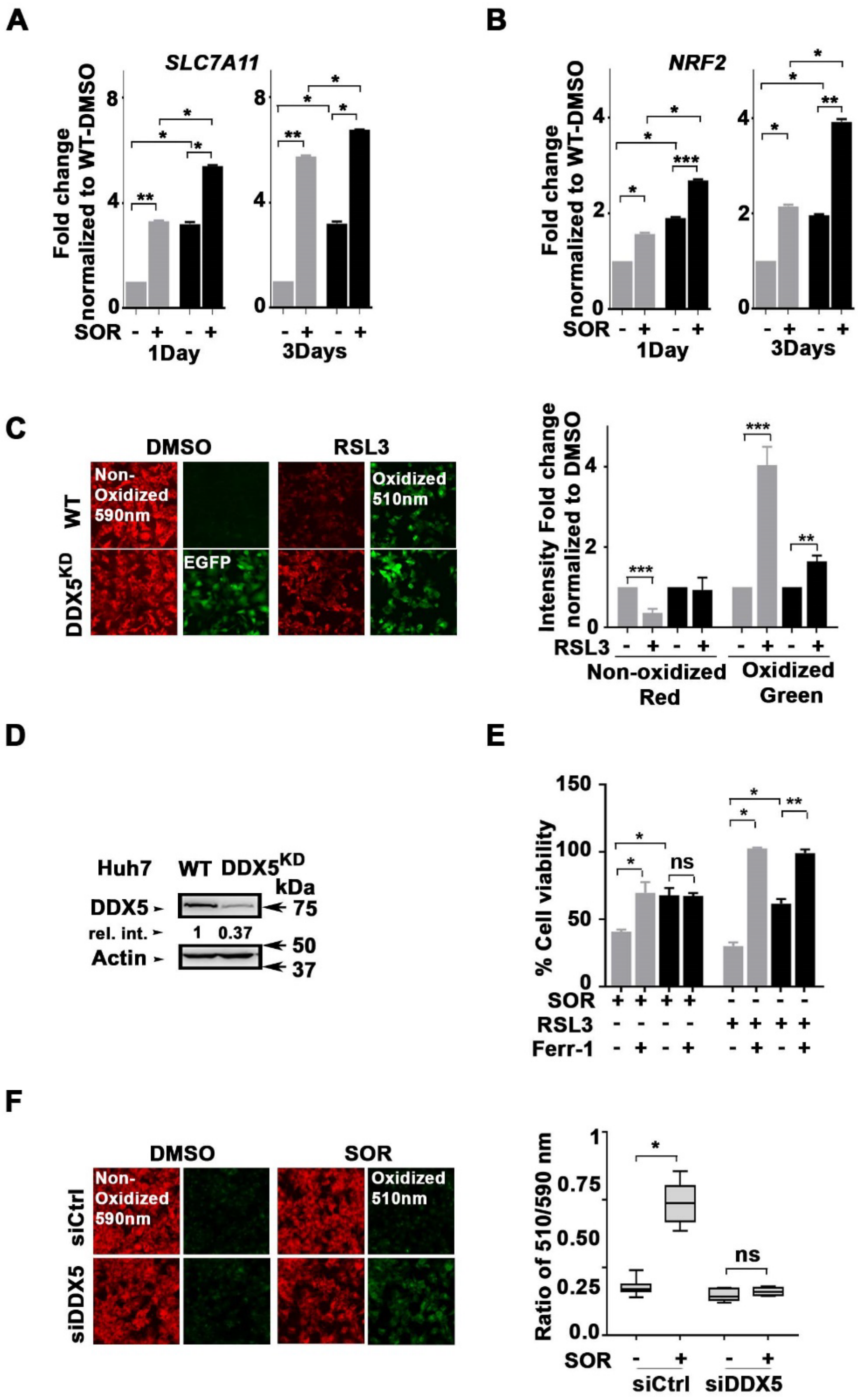


**Figure S1**. qRT-PCR of **(A)** *SLC7A11* and **(B)** *NRF2* using RNA from HepAD38$^{WT}$ and HepAD38-DDX5$^{KD}$, treated with SOR (10 µM for 1 day and 7.5 µM for 3 days, n=3). *p<0.05, ** p<0.01, ***p<0.001. **(C)** C11-BODIPY assay of HepAD38$^{WT}$ and HepAD38-DDX5$^{KD}$ treated with SOR (15 µM for 1 day). (Right panel) Quantification by ImageJ software from three independent experiments.

**(D)** Immunoblots of DDX5 using Huh7-WT and Huh7-DDX5$^{KD}$ cell lysates. **(E)** Cell viability assays using Huh7-WT and Huh7-DDX5$^{KD}$ cells treated as indicated for 24 h with 10 µM SOR, 0.5 µM RLS3 +/- 10 µM ferrostatin-1 (Ferr-1). n=3 *p<0.05, ** p<0.01, ***p<0.00. **(F)** C11-BODIPY assays of Huh7-WT cells transfected with 50 pM siCtrl or siDDX5 and treated with SOR for 24h. (Right panel) Quantification of 510nm/590 ratio from three independent experiments.

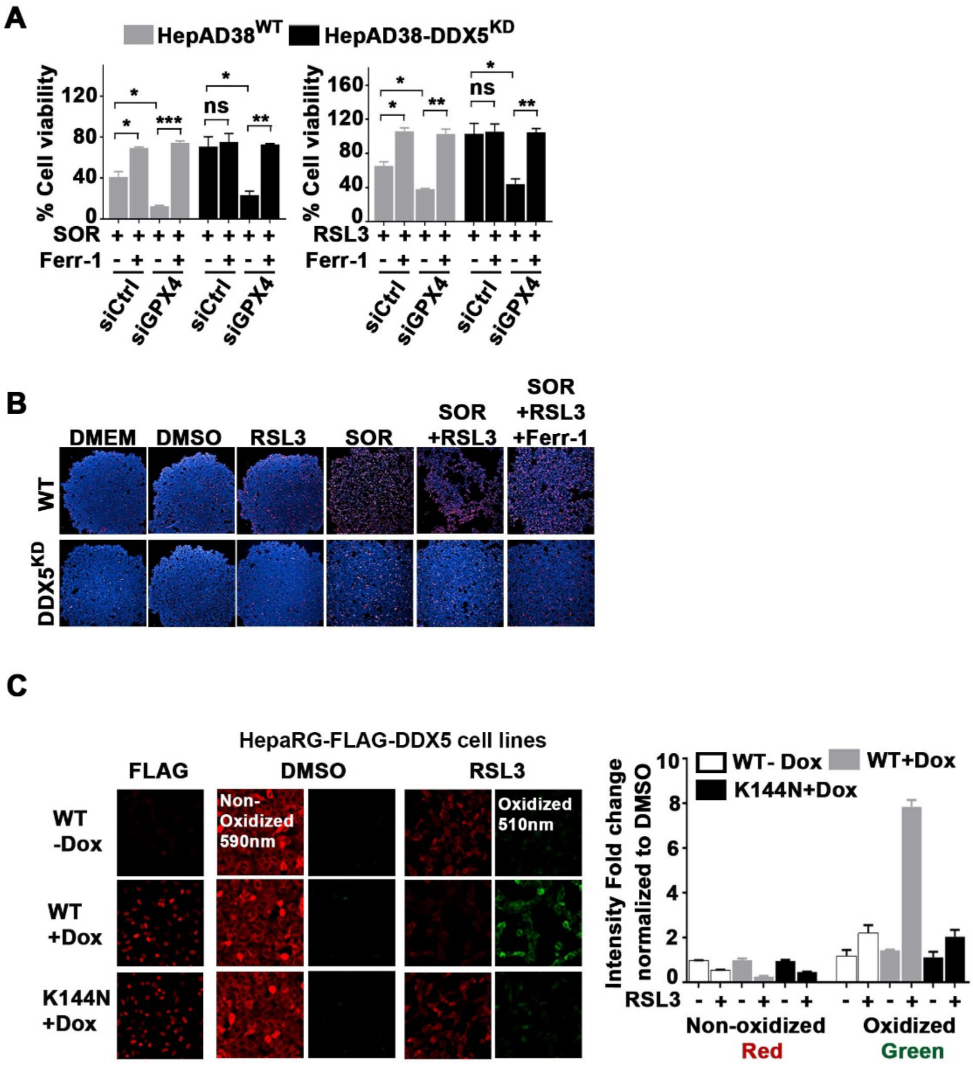


**Figure S2. (A)** Cell viability assays of HepAD38$^{WT}$ and HepAD38-DDX5$^{KD}$ cells treated as indicated for 24 h with 10 µM SOR, or 0.5 µM RLS3 +/- 10 µM ferrostatin-1 (Ferr-1), and transfected with 50 pM siRNAs (Ctrl or GPX4) for 48h. n=3 *p<0.05, ** p<0.01, ***p<0.00 **(B)** 3-D spheroid assays of HepAD38$^{WT}$ and HepAD38DDX5$^{KD}$ cells grown with indicated inhibitors (10 µM SOR, 0.5 µM RLS3 +/- 10 µM Ferr-1) for 24h. Blue is DAPI, Red is ethidium homodimer-1, staining dead cells. **(C)** Doxycycline-inducible HepaRG cell lines treated with doxycycline for 24 h to express WT FLAG-DDX5 or inactive FLAG-K144N-DDX5. FLAG immunofluorescence shows FLAG-DDX5 expression by doxycycline addition. C11-BODIPY$^{581/591}$ fluorescence microscopy of inducible HepaRG cell lines, treated with 2.0 µM RSL3 for 24 h. (Right panel) Quantification by ImageJ software of the 510nm/590 ratio.

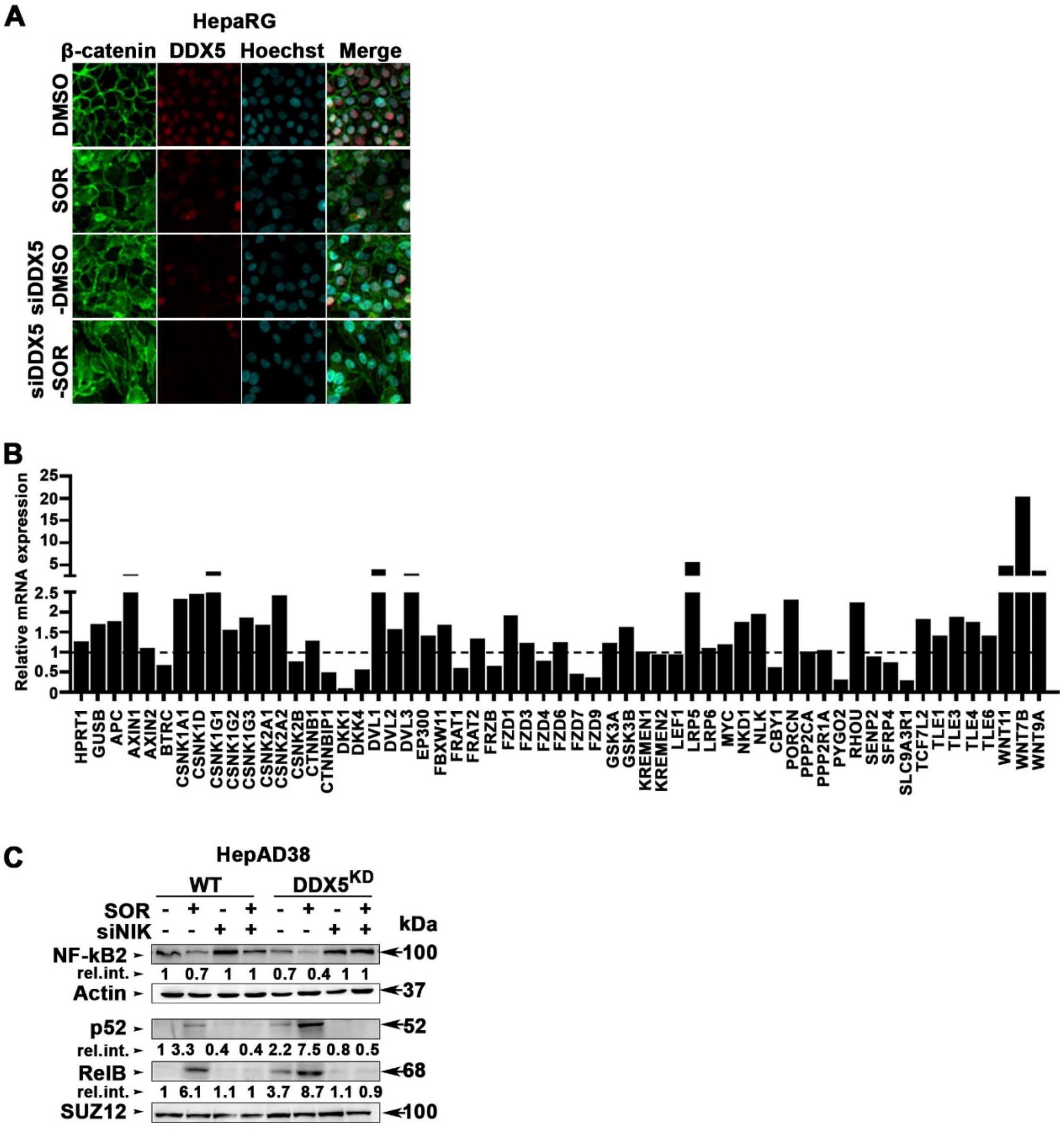


**Figure S3: (A)** Immunofluoresence microscopy of β-catenin and DDX5 in HepaRG cells treated with SOR (7.5 µM for 2 days) or siDDX5 (50 pM), as indicated. **(B)** Quantification of Wnt signaling genes in HepAD38 cells +/- SOR (7.5 µM for 3 days) using a Wnt microarray from Thermofisher. Results shown represent the average of two independent RNA isolations. **(C)** Immunoblots with indicated antibodies, using cytoplasmic (Actin loading control) and nuclear (SUZ12 loading control) lysates from indicated cell lines transfected with indicated siRNAs (50 pM): siNIK (+) or siCtrl (-) for 72 h and treated with 7.5 µM SOR (+) or DMSO (-) for the last 48 h. A representative immunoblot shown from three independent experiments

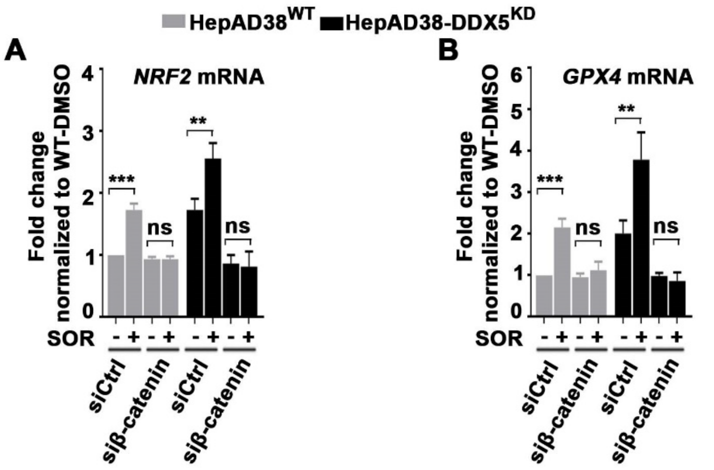


**Figure S4.** RT-PCR quantification of *NRF2* **(A)** and *GPX4* **(B)**, using RNA from HepAD38$^{WT}$ and HepAD38-DDX5$^{KD}$ cells treated +/- SOR (7.5 µM for 3 days) and transfected with siCrtl or siβ-catenin siRNAs (50 pM). n=3 *p<0.05, ** p<0.01.

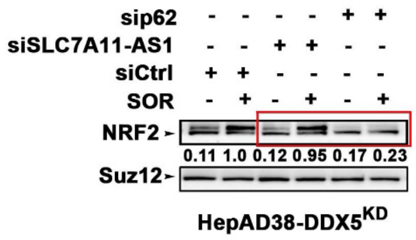


**Figure S5.** HepAD38-DDX5$^{KD}$ cells transfected with 50 pM of siRNAs: siCtrl, sip62/SQSTM1 & siSLC7A11-AS1; 24 hours later, cells were treated with SOR (7.5µM for 2 days). Nuclear extracts were immunobloted with indicated antibodies.

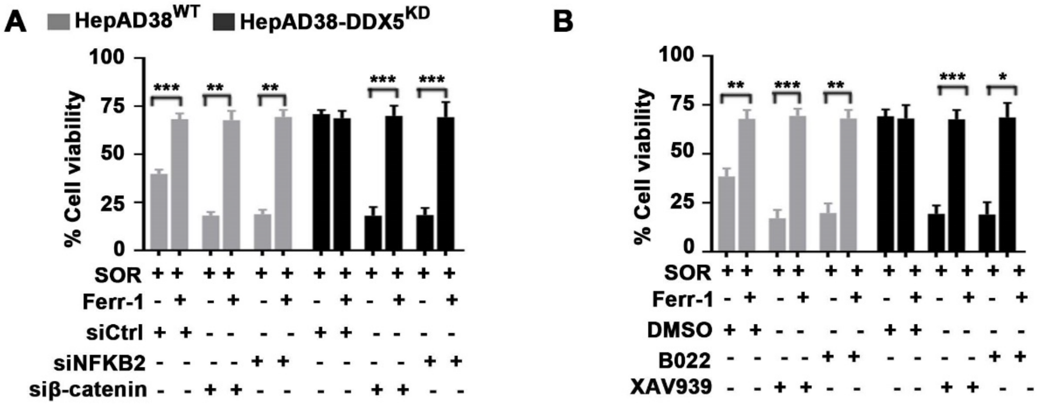


**Figure S6. (A)** MTS cell viability assays of HepAD38$^{WT}$ and HepAD38-DDX5$^{KD}$ cells transfected with 50 pM each of indicated siRNAs (siCtrl and siNFKB2), for 48 h, followed by treatment with 10 µM SOR, +/- 10 µM ferrostatin-1 (Ferr-1) for an additional 24 h, or combination with +B022 (5.0 µM) or XAV939 (20 µM) in **(B).** n=3 ***p<0.001.

**Supplementary Materials and Methods**

**Sphere Assays**: HepAD38$^{WT}$ and HepAD38-DDX5$^{KD}$ cells (1x10$^{3}$) seeded in ultra-low attachment 96-well plates (Corning spheroid microplates) were treated with sorafenib (10 μM), RSL3 (0.5μM) or DMSO for 24 h. Cells stained with NucRed Dead 647 ReadyProbes Reagent and Hoechest33342, imaged by Opera Phoenix.

**Immunoblotting:** Cells were lysed (15 min, 4$^{o}$C) in lysis buffer (Cell Signaling), sonicated on ice for 30 sec and clarified by centrifugation (13,000 rpm, 15 min, 4$^{o}$ C). Protein concentration was determined using BCA assay. All samples were diluted to 1μg/μL using 4x dye (Biorad) and equal amounts of proteins (5.0 – 40 μg per lane) were run on SDS-PAGE. Following electrophoresis, proteins transferred to nitrocellulose membrane via wet transfer (200 mA, 45 - 90 min at 4$^{o}$ C). Following transfer, membranes were blocked with 3% (w/v) BSA in Tris-buffered saline containing 0.1% (v/v) Tween 20 (TBST) and incubated with primary antibody in 3% (w/v) BSA in TBST for 1hr at room temperature, followed by incubation in secondary antibody (1:2000 dilution) in 3% BSA in TBST for 1h at room temperature. Three washes performed after primary and secondary antibody incubations. Protein bands detected by chemiluminescence using Pierce ECL (Biorad). Densitometric analysis of immunoblots performed using ImageJ software. Immunoblots are representative of three independent experiments. Antibodies used are listed in Supplementary Table S2:

**Immunofluorescence microscopy:** HepaRG cells (1x10$^{5}$) seeded on coverslips in 12-well plates and incubated at 37$^{o}$C overnight. siRNAs (50 pM) were transfected using RNAimax for 24h, followed by 24h treatment with sorafenib; next, cells were fixed using 4% PFA for 10 min at room temperature. Following fixation, cells were permeabilized with Phosphate Buffered Saline (PBS) containing 0.1% (v/v) Triton X-100 for 15 min at room temperature. Blocking used 10% goat serum in PBS containing

0.1% (v/v) Tween-20 (PBST) for 1 h at room temperature. Primary antibody to detect beta-catenin, conjugated to fluorophore, incubated in blocking solution for 1 hr at room temperature; primary antibody to detect DDX5 incubated at $4^0$C overnight, followed by secondary antibody conjugated to fluorophore in blocking solution, and incubated for 45 min at room temperature. Three washes (5 min each) performed after primary and secondary antibody incubations. One min incubation with Hoechst diluted in PBST to a final concentration of 2.0 μg/mL to stain DNA. Microscopy performed using an Olympus Fluoview confocal microscope. Antibodies used listed in Supplementary Table S2.

**RNA preparation and qRT-PCR:** RNA was isolated using Purelink mRNA Mini kit (Life Technologies) or Direct Zol RNA miniprep kit (Zymo Research). cDNA synthesized from 1.0 μg total RNA using iSCRIPT cDNA synthesis kit (Biorad). qRT-PCR performed using SYBR green (Roche) in triplicates, normalized to GAPDH.

**Supplementary Table S1: List of Plasmids and siRNAs**

| **Plasmids, siRNAs** | **Source** |
|---|---|
| Renilla luciferase vector | Addgene (#27163) |
| TOPFlash vector | Addgene (#12456) |
| NanoLuc® Reporter Vector with NF-κB Response Element | Promega( # N1111) |
| MAP3K14-Luciferase vector | Kindly provided by Dr. Inoue, J. (ref. 47) |
| siCtrl | ThermoFisher Scientific (#4390843) |
| siDDX5-1 | ThermoFisher Scientific (#4392420, assay id s4007) |
| siDDX5-2 | ThermoFisher Scientific (#4392420, assay id s4008) |
| siNRF2 | ThermoFisher Scientific (#107966, assay id AM16708) |
| siGPX4 | ThermoFisher Scientific (#10848, assay id AM16708) |
| siNIK | ThermoFisher Scientific (#110821, assay id AM16708) |
| siCTNNB1(catenin beta 1) | ThermoFisher Scientific (#146154, assay id AM16708) |
| siNFKB2 | ThermoFisher Scientific (#106835, assay id AM16708) |
| siSQSTM1/p62 | Horizon (SMARTpool, L-010230-00-0005) |

**Supplementary Table S2: Antibodies Used**

| **Antibody** | **Dilution** | **Application** | **Source** |
|---|---|---|---|
| Mouse α-Human p68 | 1:1000 in 2% BSA in TBST | Western Blot | Millipore Sigma (#05-850) |
| Rabbit α-Human DDX5 | 1:1000 in 2% BSA in TBST | Western Blot | Cell Signaling Technologies (#14994S) |
| Mouse α-Human keap1 | 1:1000 in 2% BSA in TBST | Western Blot | Santa Cruz Biotechnology (#sc-365626) |
| Rabbit α-Human SQSTM1/p62 | 1:1000 in 2% BSA in TBST | Western Blot | Cell Signaling Technologies (#5114S) |
| Rabbit α-Human NRF2 | 1:1000 in 2% BSA in TBST | Western Blot | Cell Signaling Technologies (#12721S) |
| Rabbit α-Human NF-κB2 p100/p52 | 1:1000 in 2% BSA in TBST | Western Blot | Cell Signaling Technologies (#4882S) |
| Mouse α-Human RelB | 1:1000 in 2% BSA in TBST | Western Blot | Santa Cruz Biotechnology (#sc-48366) |
| Mouse α-Human NFκB p65 | 1:1000 in 2% BSA in TBST | Western Blot | Santa Cruz Biotechnology (#sc-8008) |
| Rabbit α-Human Suz12 | 1:1000 in 2% BSA in TBST | Western Blot | Cell Signaling Technologies (#3737S) |
| Mouse α-Human Actin | 1:1000 in 2% BSA in TBST | Western Blot | Sigma (#A5441) |
| Horse α-Mouse secondary | 1:2000 in 2% BSA in TBST | Western Blot | Vector Laboratories (#PI-2000) |
| Goat α-Rabbit secondary | 1:2000 in 2% BSA in TBST | Western Blot | Vector Laboratories (#PI-1000) |
| Mouse FLAG | 1:1000 in 2% BSA in TBST | Western Blot | Sigma (#F1804) |
| Rabbit α-Human DDX5 | 1:100 in 10% Goat serum in PBST | IF | Cell Signaling Technologies (#9877S) |
| Mouse α-Human β-catenin | 1:100 in 10% Goat serum in PBST | IF | Cell Signaling Technologies (#2677S) |
| Goat α- Mouse Alexa Fluor 546 | 1:400 in 10% Goat serum in PBST | IF | ThermoFisher Scientific (#A-11030) |
| Goat α- Rabbit Alexa Fluor 546 | 1:400 in 10% Goat serum in PBST | IF | ThermoFisher Scientific (#A-11010) |

**<u>Supplementary Table S3</u>: Primer sequences**

| **Primer** | **5' – Sequence – 3'** |
|---|---|
| DDX5-F | AGCAAGTGAGCGACCTTATC |
| DDX5-R | CATCCTTCATGCCTCCTCTAC |
| GAPDH-F | CCCTTCATTGACCTCAACTACA |
| GAPDH-R | ATGACAAGCTTCCCGTTCTC |
| UBc-F | CCTGGAGGAGAAGAGGAAAGAGA |
| UBc-R | TTGAGGACCTCTGTGTATTTGTCA |
| NRF2-F | TTCAGATGCCACAGTCAACAC |
| NRF2-R | GGCATGCTGTTGCTGATACT |
| SLC7A11-F | TCCTGCTTTGGCTCCATGAACG |
| SLC7A11-R | AGAGGAGTGTGCTTGCGGACAT |
| GPX4-F | GCCTTCCCGTGTAACCAGT |
| GPX4-R | GCGAACTCTTTGATCTCTTCGT |
| NIK-F | TTGGTTGGGGAGATCGGCGCT TG |
| NIK-R | GGGGCTGAACTCTTGGCTATTCTC |

**Supplementary Table S4: Reagents, Chemical inhibitors, and Kits**

| **Reagents, Chemical inhibitors, Kits** | **Source** |
|---|---|
| Cycloheximide (CHX) | Selleck Chemicals (#S7418) |
| MG132 | Selleck Chemicals (#S2619) |
| Sorafenib | Selleck Chemicals (#S7397) |
| ICG-001 | Selleck Chemicals (#S2662) |
| XAV-939 | Selleck Chemicals (#S1180) |
| RSL3 | Selleck Chemicals (#S8155) |
| Lenvatinib (E7080) | Selleck Chemicals (#S1164) |
| Regorafenib (BAY 73-4506) | Selleck Chemicals (#S1178) |
| CellTiter 96® AQueous One Solution Cell Proliferation Assay (MTS) | Promega (#G3580) |
| PCR Mycoplasma Detection Kit | Abm (#G238) |
| Dual-Luciferase® Reporter Assay System | Promega (#1980) |
| Cell Lysis Buffer (10X) | Cell Signaling Technology (#9803) |
| LightCycler® 480 SYBR Green I Master | Roche (#04887352001) |
| iScript™ cDNA Synthesis Kit | Biorad (#1708891) |
| Nitrocellulose Membrane, Roll, 0.2 µm | Biorad (#1620112) |
| LightCycler® 480 Sealing Foil | Roche (#04729757001) |
| LightCycler® 8-Tube Strips (white) | Roche (#06612601001) |
| DMSO | Sigma (#D8418-50ML) |

| | |
|---|---|
| Tween™ 20 | ThermoFisher Scientific (BP337-500) |
| Triton™ X-100 | Sigma (#T8787-100ML) |
| Costar® 6-well Ultra-Low Attachment Plates | Corning (#3471) |
| Tetracycline hydrochloride | Sigma (#T7660-5G) |
| Bovine Serum Albumin | Sigma (#A9647-100G) |
| Pierce™ ECL Western Blotting Substrate | ThermoFisher Scientific (#32106) |
| Geneticin™ Selective Antibiotic (G418 Sulfate) | ThermoFisher Scientific (#10131027) |
| Pierce™ BCA Protein Assay Kit | ThermoFisher Scientific (#23227) |
| Lipofectamine™ 3000 Transfection Reagent | ThermoFisher Scientific (#L3000015) |
| Lipofectamine™ RNAiMAX Transfection Reagent | ThermoFisher Scientific (#13778150) |
| Restore™ PLUS Western Blot Stripping Buffer | ThermoFisher Scientific (#46430) |
| RNeasy Mini Kit | Qiagen (#74104) |
| Hoechst 33342 Solution (20 mM) | ThermoFisher Scientific (#62249) |
| Corning® 96 Well Black Polystyrene Microplate | Corning(#CLS3603-48EA) |
| CellTiter-Glo® 2.0 Cell Viability Assay | Promega( #G9242) |
| BODIPY™ 581/591 C11 (Lipid Peroxidation Sensor) | ThermoFisher Scientific ( # D3861) |
| 29 mm Glass bottom dish with 14 mm micro-well #1.5 cover glass | NC0662883（D29-14-1.5-N） |
| Nano-Glo® Luciferase Assay System | Promega( # N1110) |
| RNAlater™ Stabilization Solution | ThermoFisher Scientific ( # AM7024) |
| PEG400 | Selleck Chemicals (#S6705) |

| | |
|---|---|
| NE-PER™ Nuclear and Cytoplasmic Extraction Reagents | ThermoFisher Scientific ( # 78835) |
| Pierce™ 16% Formaldehyde (w/v), Methanol-free | ThermoFisher Scientific ( # 28908) |
| B022 | MedChemExpress( # HY-120501) |
| Protease Inhibitor Cocktail (100X) | Cell Signaling Technology (#5871) |
| NucRed™ Dead 647 ReadyProbes™ Reagent (TO-PRO-3 iodide) | ThermoFisher Scientific ( # R37113) |
